# Reduced Suprathreshold Auditory Nerve Responses are Associated with Thinner Temporal Cortex and Slower Processing Speed in Presbycusis

**DOI:** 10.1101/2020.02.12.945337

**Authors:** Paul H. Delano, Chama Belkhiria, Rodrigo C. Vergara, Melissa Martínez, Alexis Leiva, Maricarmen Andrade, Bruno Marcenaro, Mariela Torrente, Juan Cristóbal Maass, Carolina Delgado

## Abstract

Epidemiological evidence shows an association between hearing loss and dementia in elderly people. However, the mechanisms that connect hearing impairments and cognitive decline are still unknown. Here we propose that a suprathreshold auditory-nerve impairment is associated with cognitive decline and brain atrophy. Methods: audiological, neuropsychological, and brain structural 3-Tesla MRI data were obtained from elders with different levels of hearing loss recruited in the ANDES cohort. The amplitude of waves I (auditory nerve) and V (midbrain) from auditory brainstem responses were measured at 80 dB nHL. We also calculated the ratio between wave V and I as a proxy of a suprathreshold brainstem function. Results: we included a total of 101 subjects (age: 73.5 ± 5.2 years (mean ± SD), mean education: 9.5 ± 4.2 years, and mean audiogram thresholds (0.5-4 kHz): 25.5 ± 12.0 dB HL). We obtained reliable suprathreshold waves V in all subjects (n=101), while replicable waves I were obtained in 92 subjects (91.1%). Partial Spearman correlations (corrected by age, gender, education and hearing thresholds) showed that reduced suprathreshold wave I responses were associated with thinner bilateral medial and inferior temporal cortex and, with slower processing speed as evidenced by the Trail-Making Test-A and digit symbol tests. Non-significant correlations were obtained between wave I amplitudes and other cognitive domains. Conclusions: These results evidence that reduced suprathreshold auditory nerve responses in presbycusis are associated with slower processing speed and brain structural changes in the temporal cortex.

## Introduction

Epidemiological studies have associated hearing loss with cognitive decline in adults older than 55 years, showing that individuals with audiometric thresholds worse than 40 dB are more likely to develop dementia [1-4]. However, the mechanisms that connect this epidemiological association are still under research [5]. Age-related hearing loss or presbycusis is characterized by bilateral hearing loss, degraded speech understanding, and impaired music perception, especially in background noise conditions [6,7]. Presbycusis is also associated with executive dysfunction [8,9] and with brain atrophy in the temporal lobe [10,11]. Moreover, recent studies in presbycusis have shown cortical atrophy in regions beyond the auditory cortex, including the cingulate cortex and parietal regions [9,12].

In addition to audiogram threshold elevations, hearing impairments in presbycusis can also be due to an altered suprathreshold function [13]. In rodents, suprathreshold brainstem responses have been extensively studied in models of acoustic injury, in which after a transient acoustic trauma, there is a temporary auditory threshold elevation that recovers completely, but a permanent reduction in the amplitude of auditory nerve responses is observed at supra-thresholds levels [14,15]. In humans, the reduction of the amplitude of wave I from auditory brainstem responses (ABR) without alterations in auditory thresholds and otoacoustic emissions levels has been termed as hidden hearing loss (HHL) [16]. The underlying structural abnormality found in animals with HHL is the loss of synapses between inner hair cells and auditory nerve neurons, a histologic feature that has been termed as cochlear synaptopathy [14,17,18]. Importantly, evidence in animals shows that cochlear synaptopathy is a contributor of the early pathophysiological process of presbycusis [19].

In humans, the suprathreshold amplitude of ABR wave I has been reported to be reduced in patients with tinnitus and normal audiograms [16], and in subjects exposed to noise [20], suggesting that HHL might be part of the pathophysiological mechanisms of these conditions. In addition, HHL has been proposed as one of the mechanisms that might degrade speech perception in noisy environments [21]. In this line, a reduction in the amplitude of suprathreshold auditory nerve responses could be considered as an early stage of hearing impairment, which can be detected before hearing loss becomes clinically evident. Whether these suprathreshold abnormalities are associated with cognitive impairment and structural brain changes in humans is unknown. Here, we hypothesize that a reduction in the amplitude of supra-thresholds auditory-nerve responses (ABR wave I) is associated with brain atrophy and cognitive decline in the elderly.

## Methods

### Subjects

The ANDES (Auditory and Dementia study) project is a prospective cohort of non-demented Chilean elders (≥65 years) with a Mini-Mental State Examination (MMSE) > 24, with different levels of age-related hearing impairment and no previous use of hearing aids. Inclusion criteria were: preserved functionality measured by the Pfeffer activities questionnaire [22], auditory brainstem responses evaluated at 80 dB nHL, and magnetic resonance imaging (MRI) at 3 Tesla. Exclusion criteria for recruitment were: (i) other causes of hearing loss different from presbycusis; (ii) previous use of hearing aids (iii); stroke or other neurological disorders; (iv) dementia; and (v) major psychiatric disorders. All procedures were approved by the Ethics Committee of the Clinical Hospital of the University of Chile, protocol number: OAIC 752/15. All subjects gave written informed consent in accordance with the Declaration of Helsinki.

### Auditory evaluations

Hearing impairments were evaluated with threshold and supra-threshold tests. All auditory evaluations were assessed inside a sound attenuating room and were obtained by an experienced audiologist who was blind to cognitive and MRI evaluations. We obtained audiometric thresholds using a calibrated audiometer (AC40e, Interacoustics®) for each ear at 0.125, 0.250, 0.5, 1, 2, 3, 4, 6 and 8 kHz. Pure tone averages (PTA) were computed for each ear using 0.5, 1, 2 and 4 kHz thresholds. The better hearing ear was used for analyses. Distortion product otoacoustic emissions (DPOAE) (2f1-f2) were elicited using eight pairs of primary tones (f1 and f2) with f2/f1 ratio=1.22, and delivered at 65 and 55 dB SPL (ER10C, Etymotic Research®). DPOAE were measured at eight different frequencies per ear, between 707 and 3563 Hz. For subsequent analyses we counted the number of detected DPOAE, a value that considering both ears, goes from 0 to 16 (see [9] for more details on DPOAE analysis). ABR waveforms were averaged with alternating clicks presented at supra-thresholds levels (2000 repetitions, 80 dB nHL, bandpass 0.1-3 kHz, stimulus rate 21.1 Hz, EP25, Eclipse, Interacoustics®). The amplitudes of waves I and V were measured from peak to trough, and wave latencies from peaks. For computing wave V/I ratios, in those cases with no measurable wave I (n=9, see results section), the minimum amplitude value that we obtained for wave I (0.02 μV) was used.

### Neuropsychological assessment

Subjects and their relatives were evaluated by a neurologist with a complete structured medical, functional and cognitive interview. Cognitive performance was assessed by an experienced psychologist in cognitive tests, including the MMSE adapted for the Chilean population for global cognition [22,23]; the Frontal Assessment Battery (FAB), perseverative errors from the Wisconsin Card Sorting (WCS) and Trail Making Test B (TMT-B) for measuring executive function [24]; the Trail Making Test A (TMT-A) and digit symbol for processing speed [25]; the Boston Nominating Test for Language [26]; the Rey-Osterrieth Complex Figure Test for Visuospatial Abilities [27]; and the free recall of the Free and Cued Selective Reminding Test (FCSRT) to explore verbal episodic memory [28,29].

### Magnetic resonance imaging

Neuroimaging data were acquired by a MAGNETOM Skyra 3-Tesla whole-body MRI Scanner (Siemens Healthcare GmbH®, Erlangen, Germany) equipped with a head volume coil. T1-weighted magnetization-prepared rapid gradient echo (T1-MPRAGE) axial images were collected, and parameters were as follows: time repetition (TR) = 2300 ms, time echo (TE) = 232 ms, matrix = 256 × 256, flip angle = 8°, 26 slices, and voxel size = 0.94 × 0.94 × 0.9 mm3. T2-weighted turbo spin echo (TSE) (4500 TR ms, 92 TE ms) and fluid attenuated inversion recovery (FLAIR) (8000 TR ms, 94 TE ms, 2500 TI ms) were also collected to inspect structural abnormalities. A total of 440 images were obtained during an acquisition time of 30 minutes per subject.

### Morphometric analyses

To determine the structural brain changes of controls and individuals with presbycusis, we measured the volume and thickness of bilateral cortical regions. FreeSurfer (version 6.0, http://surfer.nmr.mgh.harvard.edu) was used with a single Linux workstation using Centos 6.0 for T1-weighted images analysis of individual subjects. The FreeSurfer processing involved several stages, as follows: volume registration with the Talairach atlas, bias field correction, initial volumetric labeling, nonlinear alignment to the Talairach space, and final volume labeling. We used the “recon-all” function to generate automatic segmentations of cortical and subcortical regions. This command performs regional segmentation and processes gross regional volume in a conformed space (256×256×256 matrix, with coronal reslicing to 1 mm^3^ voxels). The function “recon-all” creates gross brain volume extents for larger-scale regions (i.e., total number of voxels per region): total grey and white matter, subcortical grey matter, brain mask volume, and estimated total intracranial volume.

Additionally, we measured the cortical thickness in native space using FreeSurfer tools. We calculated the cortical thickness of each mesh of vertices by measuring the distance between the point on one surface and the closest conforming point on the opposite surface. Then we measured the average of the two values calculated from each side to the other [30]. Based on the brain regions that have been previously studied in presbycusis [10,31] our regions of interest (ROI) were bilateral frontal, inferior, middle, superior and transverse temporal gyri, and parietal cortex. We also included as regions of interest, cortical areas that have been implicated in the neural networks of degraded speech comprehension: bilateral anterior cingulate cortex, posterior cingulate cortex (PCC), and precentral and postcentral gyri [9,11,32].

### Data analyses

Possible correlations between cognitive tests and audiological functions were evaluated by means of partial Spearman associations adjusted by age, educational level, gender and audiogram thresholds. Gender comparisons were done using Mann-Whitney tests. Comparisons between subgroups were performed with ANCOVA adjusted by age, education, audiogram thresholds and gender. This approach was maintained for two group comparisons, as t-test do not allow covariates. Bonferroni corrections were performed for multiple comparisons when comparing more than two groups. Data are shown as mean ± standard deviation. Significant differences and correlations were considered for p<0.05.

## Results

### Demographic and audiological variables

The mean age of the 101 studied subjects was 73.5 ± 5.2 years with a mean education of 9.5 ± 4.2 years, and mean PTA of the better hearing ear of 25.5 ± 12.0 dB HL. A demographic description of the 101 subjects that completed the auditory, neuropsychological, and MRI evaluations is presented in Table 1. As one of our recruitment criteria was that subjects were not using hearing aids, the majority of the enrolled individuals had normal hearing thresholds (PTA < 25 dB HL, n=55, 54.5%), while 46 subjects had some degree of hearing loss, including 33 (32.7%) with mild hearing loss (PTA ≥ 25 dB HL <40 dB HL), and 13 individuals (12.8%) with moderate hearing loss (PTA ≥ 40 dB HL) according to audiogram thresholds of the better hearing ear. Age and audiogram thresholds were significantly correlated (Spearman, rho=0.326, p=0.001), while the educational level was not correlated with PTA thresholds (Spearman, rho=0.0622, p=0.536) (Figure 1A, D).

**Table 1.**
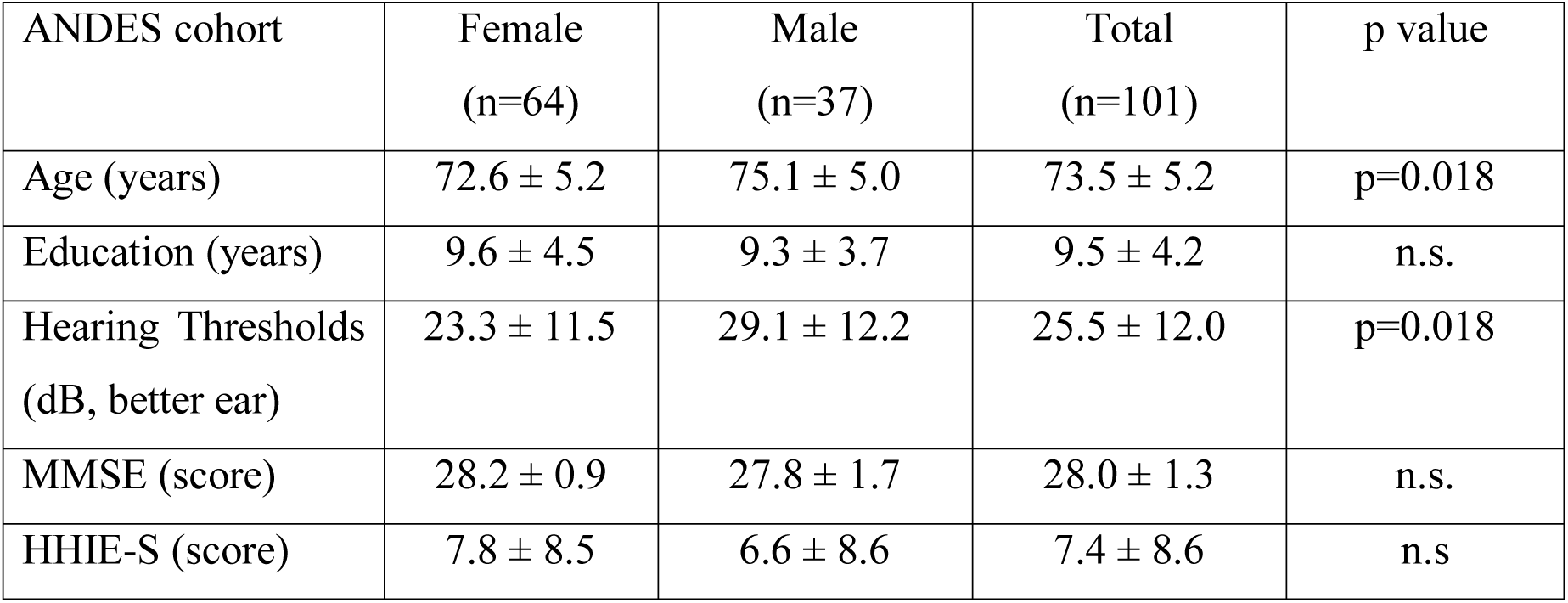
Summary of demographic data of the subjects considered in this report (obtained from ANDES cohort, n=101). Significant gender differences were obtained for age and hearing thresholds, as men are older and have worse hearing thresholds than women (p<0.05, Mann Whitney). MMSE: Mini Mental State Examination, HHIE-S: Hearing Handicap Inventory for the Elderly, ns: non-significant.

**Figure 1.**
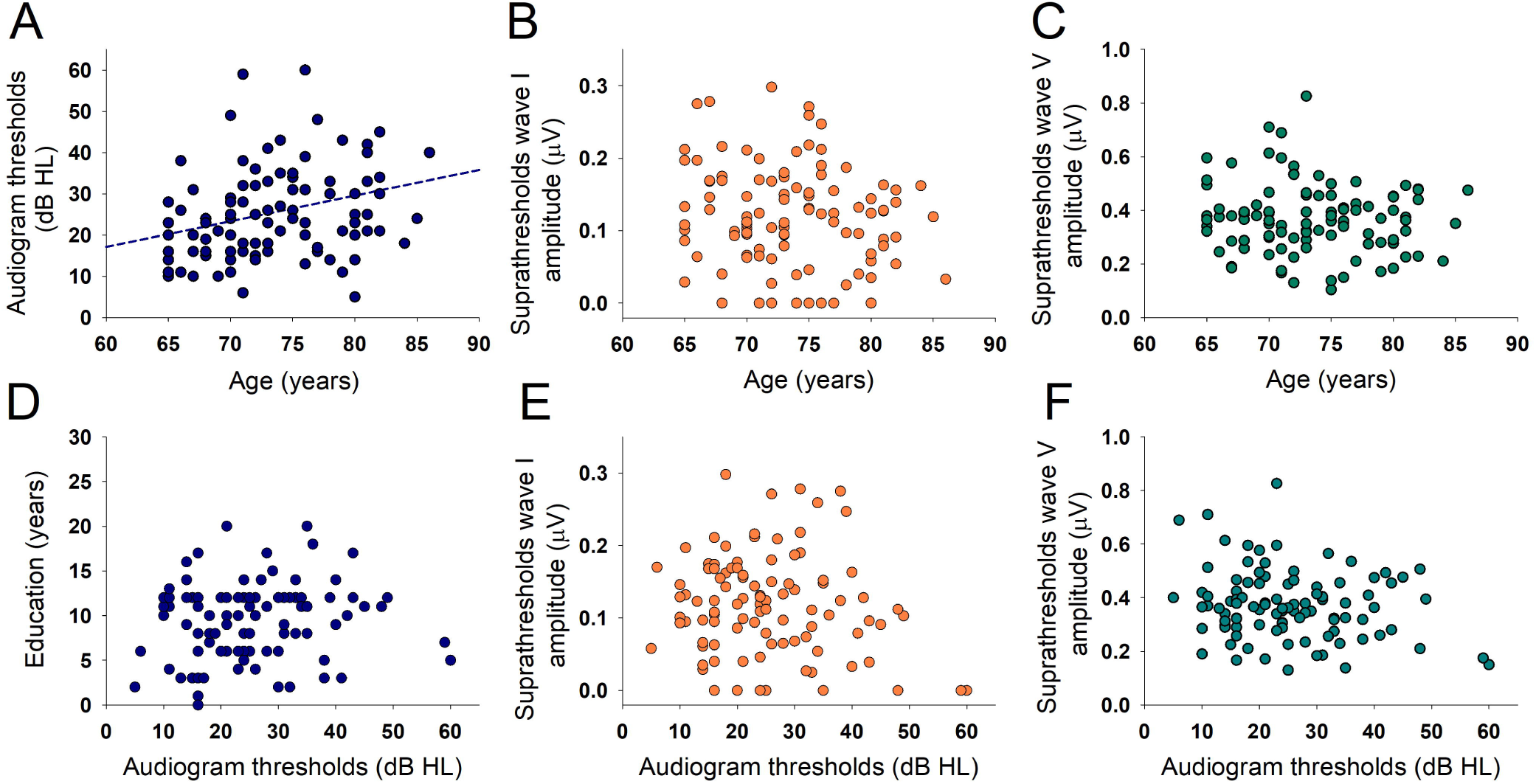
Correlations between audiogram thresholds, age, education and supra-thresholds ABR responses. A. Age and PTA were significantly correlated (Spearman, rho=0.326, p=0.001). B. and E. Scatter plots showing no correlations between the amplitude of wave I with age (in the range between 65 and 85 years) and audiogram thresholds. C. and F. Scatter plots showing no correlations between the amplitude of wave V with age (in the range between 65 and 85 years) and audiogram thresholds. D. Audiogram thresholds were not correlated with the years of education.

Regarding supra-threshold ABR responses, we obtained measurable waves V at 80 dB nHL in the 101 subjects of this study, while wave I was obtained in 92 of these subjects (91.1%). The average amplitudes of wave I and V were 0.120 ± 0.070 μV and 0.369 ± 0.129 μV respectively, while mean latencies were 5.71 ± 0.39 ms for wave V and 1.56 ± 0.14 ms for wave I. We found a significant correlation between the amplitude of wave I and wave V (Figure 2A, rho=0.323, p=0.001), while there were no correlations between the supra-threshold amplitudes of ABR waves I and V and age and audiogram thresholds (Figure 1B, C, E, F). In addition, there were non-significant differences in the amplitude of wave I when comparing subjects with hearing loss (n=46, 0.113 ± 0.79 μV) with those with normal audiogram thresholds (n=55, 0.124 ± 0.62 μV, F(1,96)=0.82, p=0.775, ANCOVA controlled for age, education and gender). Regarding suprathreshold wave V amplitudes, we also obtained non-significant effects when comparing control and hearing loss subjects (controls: n=55, 0.394 ± 0.134 μV; hearing loss; n=46, 0.340 ± 0.118 μV, F(1,96)=3.82, p= 0.054, ANCOVA controlled for age, education and gender).

**Figure 2.**
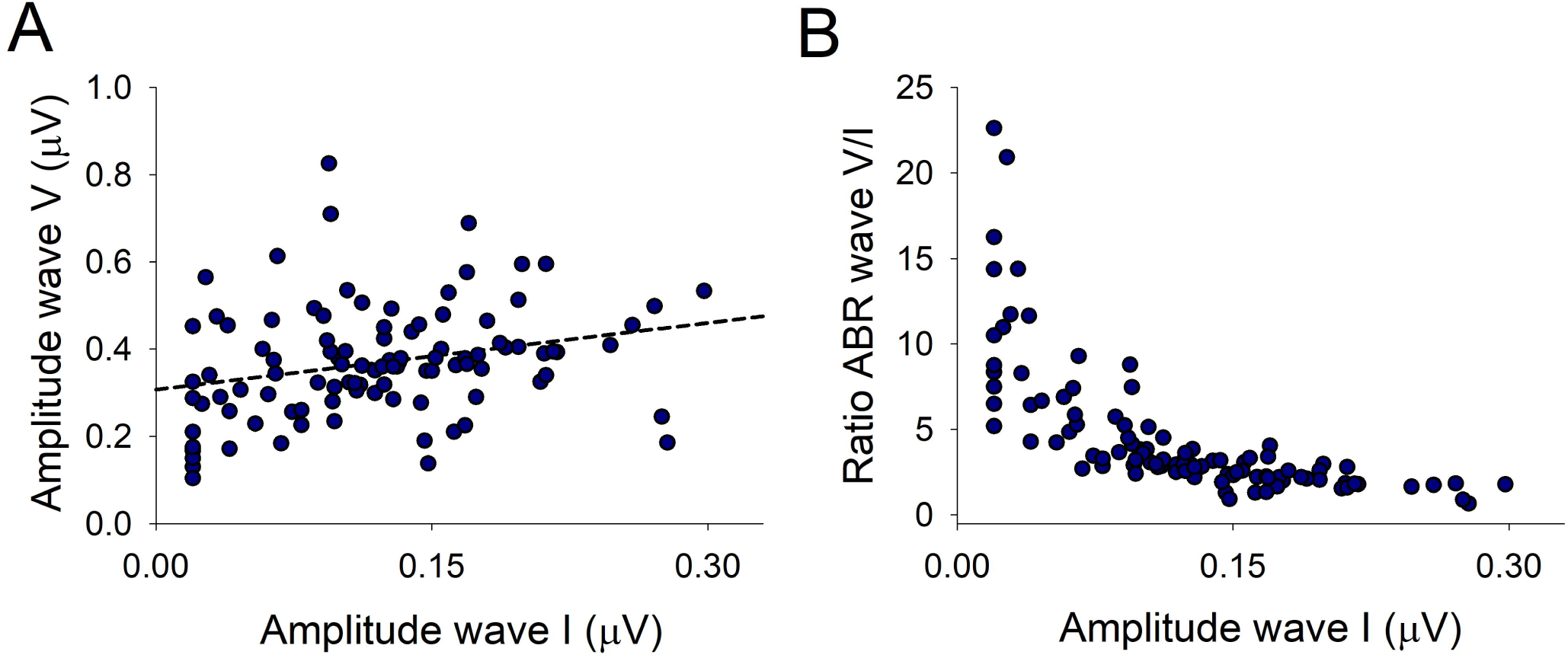
Correlations between the amplitude and ratio of suprathreshold ABR responses. A. The amplitude of wave I was significantly correlated with the amplitude of wave V (rho=0.323, p=0.001). B. Wave ABR V/I amplitude ratio plotted as a function of wave I amplitude. Notice an asymmetric distribution of wave V/I ratio as a function of wave I amplitude, showing larger wave V/I ratios for wave I amplitudes smaller than 0.15 μV.

We also calculated the ratio between waves V and I which has been used as a measure of hidden hearing loss in previous studies [16,19]. The average wave V/I ratio was 4.5 ± 3.9 (interquartile range 2.24 - 5.21). There was an asymmetric distribution of the wave V/I ratio as a function of wave I amplitude, denoting that wave V/I ratios for wave I amplitudes below 0.15 μV were significantly larger than for those above 0.15 μV (Mann-Whitney, p<0.001) (Figure 2B, Table 2). Non-significant correlations were obtained between age and audiogram thresholds with the wave V/I ratio (not shown). In addition, there were non-significant differences in the wave V/I ratio when comparing subjects with hearing loss (n=46, 4.7 ± 4.2) with those with normal audiogram thresholds (n=55, 4.4 ± 3.7, F(1,96)=0.42, p=0.519, ANCOVA controlled for age, education and gender).

**Table 2.** Demographic and neuropsychological variables compared according to the two groups with different amplitude of auditory nerve responses. ANCOVA was corrected by age, gender, education and audiogram thresholds. Note that TMT-A time is the only significant difference in cognitive performance between the groups (p<0.05*, adjusted by Bonferroni for multiple comparisons).

| ANDES cohort<br>(n=101) | Auditory nerve<br>less than<br>0.15 μV ABR wave I<br>(n=68) | Auditory nerve<br>more than<br>0.15 μV ABR wave I<br>(n=33) | p value<br>ANCOVA |
| --- | --- | --- | --- |
| Age | 74.0 ± 5.3 | 72.6 ± 4.9 | n.s. |
| Years of education | 9.7 ± 4.3 | 9.0 ± 4.1 | n.s. |
| PTA 0.5-4 kHz<br>(dB) | 26.8 ± 13.3 | 23.1 ± 8.5 | n.s. |
| DPOAE (n, both<br>ears) | 7.0 ± 5.8 | 7.8 ± 5.1 | n.s. |
| ABR wave V<br>amplitude (μV) | 0.349 ± 0.132 | 0.410 ± 0.115 | n.s. |
| Wave V/I ratio | 5.64 ± 4.32 | 2.15 ± 0.73 | p<0.001* |
| MMSE | 27.82 ± 1.40 | 28.42 ± 1.30 | n.s. |
| TMT-A (s) | 66.3 ± 31.5 | 51.9 ± 23.0 | p=0.005* |
| TMT-B (s) | 172.4 ± 84.0 | 176.8 ± 91.1 | n.s. |
| Digit symbol | 36.3 ± 14.7 | 40.5 ± 13.1 | n.s. |
| Fluency P | 10.0 ± 4.8 | 10.1 ± 4.3 | n.s. |
| Perseverative errors<br>(WCS) | 11.1 ± 9.1 | 10.2 ± 6.3 | n.s. |
| Boston nomination | 24.4 ± 3.2 | 25.0 ± 3.4 | n.s. |
| Rey Figure | 30.0 ± 5.4 | 29.4 ± 5.1 | n.s. |
| FAB | 13.3 ± 2.5 | 14.0 ± 2.0 | n.s |
| FCRST free recall | 25.9 ± 7.9 | 26.2 ± 7.0 | n.s |

As the increased wave V/I ratio might be reflecting a compensatory midbrain gain increase of wave V responses in the group with wave I < 0.15 μV, we divided data according to the amplitude of wave I into two groups: (i) those with wave I responses smaller than 0.15 μV (n=68) and (ii) those with wave I responses larger than 0.15 μV (n=33). Table 2 shows demographic, audiological and neuropsychological data comparing these two groups with different wave I amplitudes. There were no differences in age, education and hearing thresholds (assessed by audiogram and DPOAEs) between these two groups.

### Suprathreshold ABRs and cognitive assessments

Regarding cognitive tests, and after adjusting by age, education, gender, audiogram thresholds, and Bonferroni correction for multiple comparisons (10 cognitive tests), the only significant difference was obtained in the TMT-A speed, showing that the group with smaller wave I responses had slower processing speed (66.3 ± 31.5 s) than the group with larger wave I responses (51.9 ± 23.0 s, p=0.005).

Next, we performed partial Spearman correlations in the whole sample (n=101) between ABR and cognitive tests, corrected by age, education, gender and audiogram thresholds. The only cognitive tests that showed significant correlations with the amplitude of supra-threshold wave I were those that measure processing speed: the TMT-A time (Figure 3, rho= −0.27, p=0.007), and the digit symbol (rho=0.199; p=0.049), while Boston performance was inversely correlated with the latency of wave V (rho=-0.208; p=0.039) (Table 3).

**Table 3.** Partial correlations between ABR amplitudes and latencies and neuropsychological tests in the ANDES cohort (n=101). All correlations were adjusted by age, education, gender and audiogram thresholds. Notice significant correlations (shown in bold) between TMT-A time and digit symbol with the amplitude of ABR wave I. In addition, Boston was significantly correlated with the latency of wave V.

| ANDES cohort<br>(n=101) | ABR wave<br>I amplitude | ABR wave V<br>amplitude | ABR wave I<br>latency | ABR wave V<br>latency |
| --- | --- | --- | --- | --- |
| TMT-A | **rho= -0.272**<br>**p= 0.007** | rho= -0.065<br>p= 0.524 | rho=0.065<br>p=0.544 | rho=0.119<br>p=0.243 |
| Digit symbol | **rho=0.199**<br>**p=0.049** | rho=0.178<br>p=0.079 | rho=-0.079<br>p=0.461 | rho=-0.121<br>p=0.234 |
| Fluency P | rho=-0.071<br>p=0.485 | rho=-0.072<br>p=0.479 | rho=0.045<br>p=0.677 | rho=-0.119<br>p=0.243 |
| Perseverative<br>errors | rho=-0.024<br>p=0.817 | rho=-0.066<br>p=0.516 | rho=-0.051<br>p=0.634 | rho=0.135<br>p=0.186 |
| Boston<br>nomination | rho=0.068<br>p=0.504 | rho=0.161<br>p=0.114 | rho=-0.097<br>p=0.363 | **rho=-0.208**<br>**p=0.039** |
| Rey Figure | rho=-0.009<br>p=0.929 | rho=-0.110<br>p=0.285 | rho=0.038<br>p=0.724 | rho=-0.117<br>p=0.256 |
| FAB | rho=0.041<br>p=0.692 | rho=0.015<br>p=0.882 | rho=0.084<br>p=0.436 | rho=-0.062<br>p=0.542 |
| FCSRT free<br>recall | rho=-0.10<br>p=0.327 | rho=-0.120<br>p=0.238 | rho=-0.094<br>p=0.382 | rho=0.160<br>p=0.876 |

**Figure 3.**
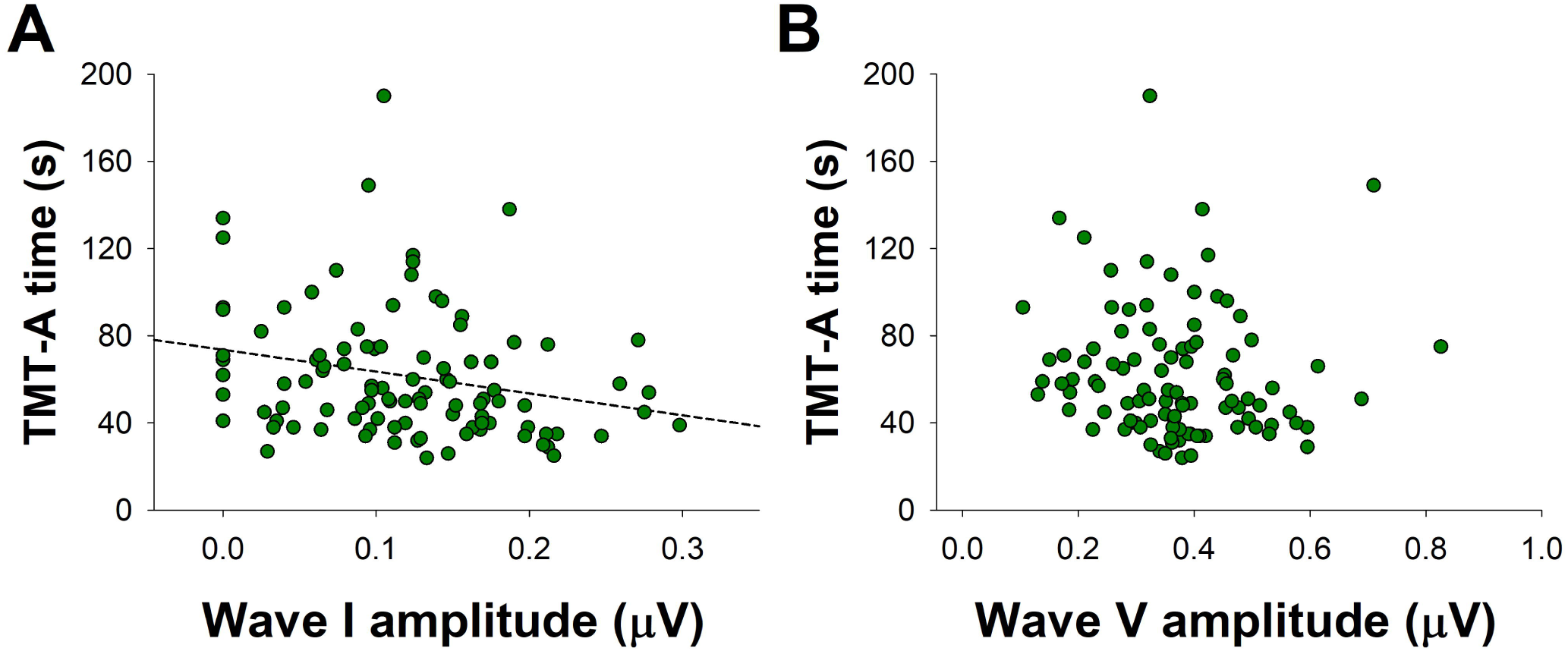
Correlations between TMT-A performance and supra-threshold ABR responses. (A) Trail-Making Test A speed is associated with the suprathreshold amplitude of wave I amplitude (rho=-0.272, p=0.007), but not with (B) the suprathreshold amplitude of wave V.

### Suprathreshold ABRs and cortical volume and thickness

We performed partial Spearman correlations between the suprathreshold amplitudes of wave I and V responses with all the cortical volumes and thickness of the ROIs in the brain (corrected by age, education, gender and audiogram thresholds). Non-significant differences were found when analyzing cortical volumes in all the ROIs between the two groups with different supra-threshold ABR amplitudes (data not shown). We found significant Spearman correlations between the amplitude of wave I and thickness of bilateral middle and inferior temporal cortex (Figure 4, Table 4). We also found significant correlations between wave I amplitude and right posterior cingulate and medial orbitofrontal cortices thickness, and for left inferior and transverse temporal cortices (Table 4). Regarding wave V, we only found a significant correlation between left inferior and transverse temporal cortices.

**Table 4.**
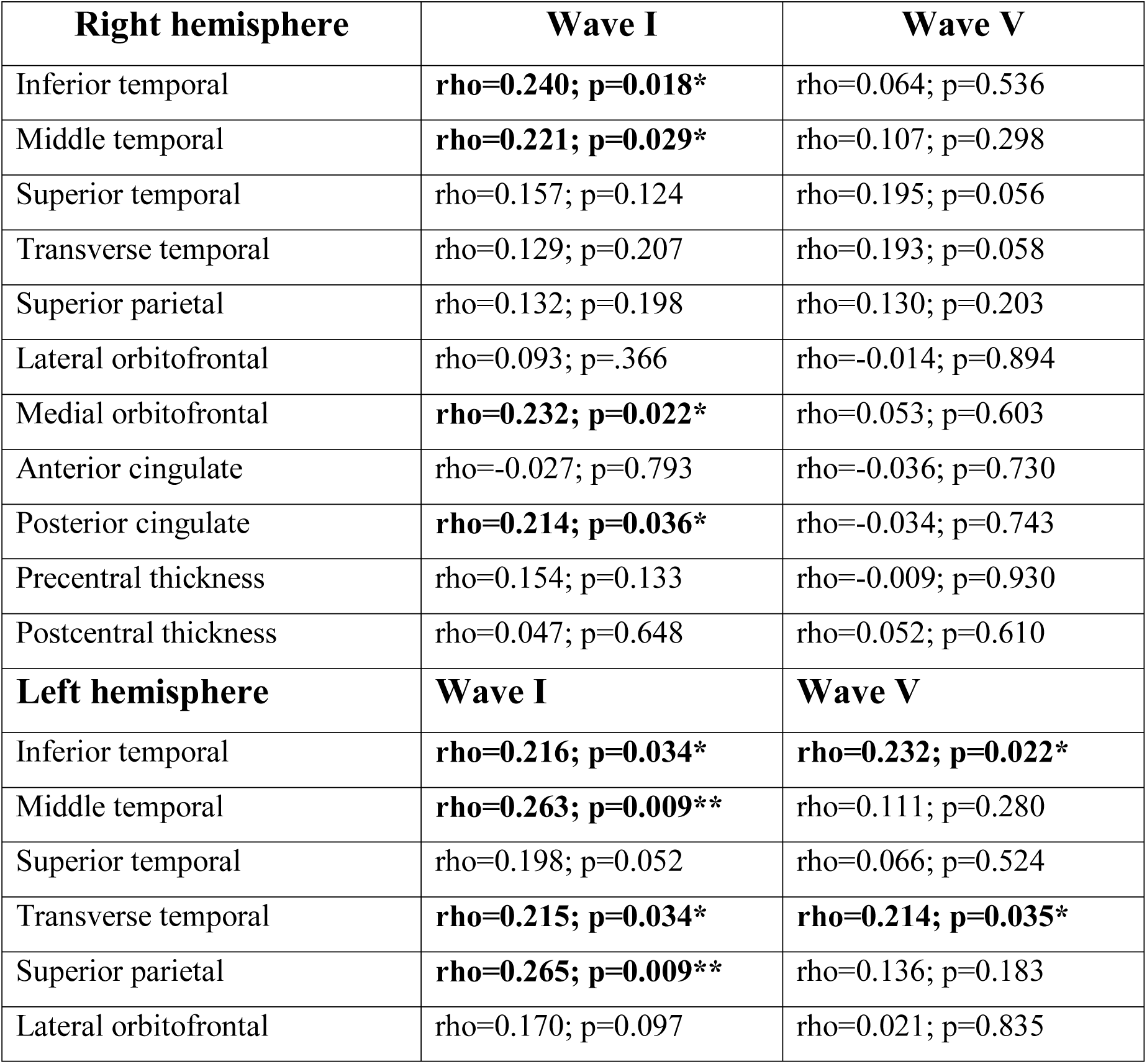
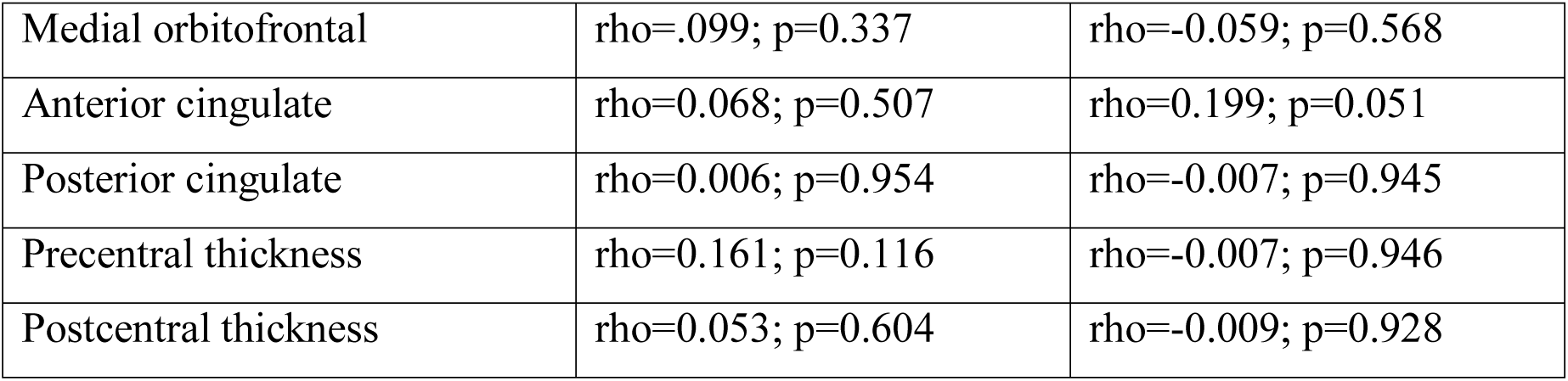
Partial correlations between ABR amplitudes and cortical thickness in presbycusis patients from the ANDES cohort (n=101). All correlations were controlled by age, education, gender and audiogram thresholds. Significant correlations are highlighted in bold font (*p<0.05; **p<0,01).

**Figure 4.**
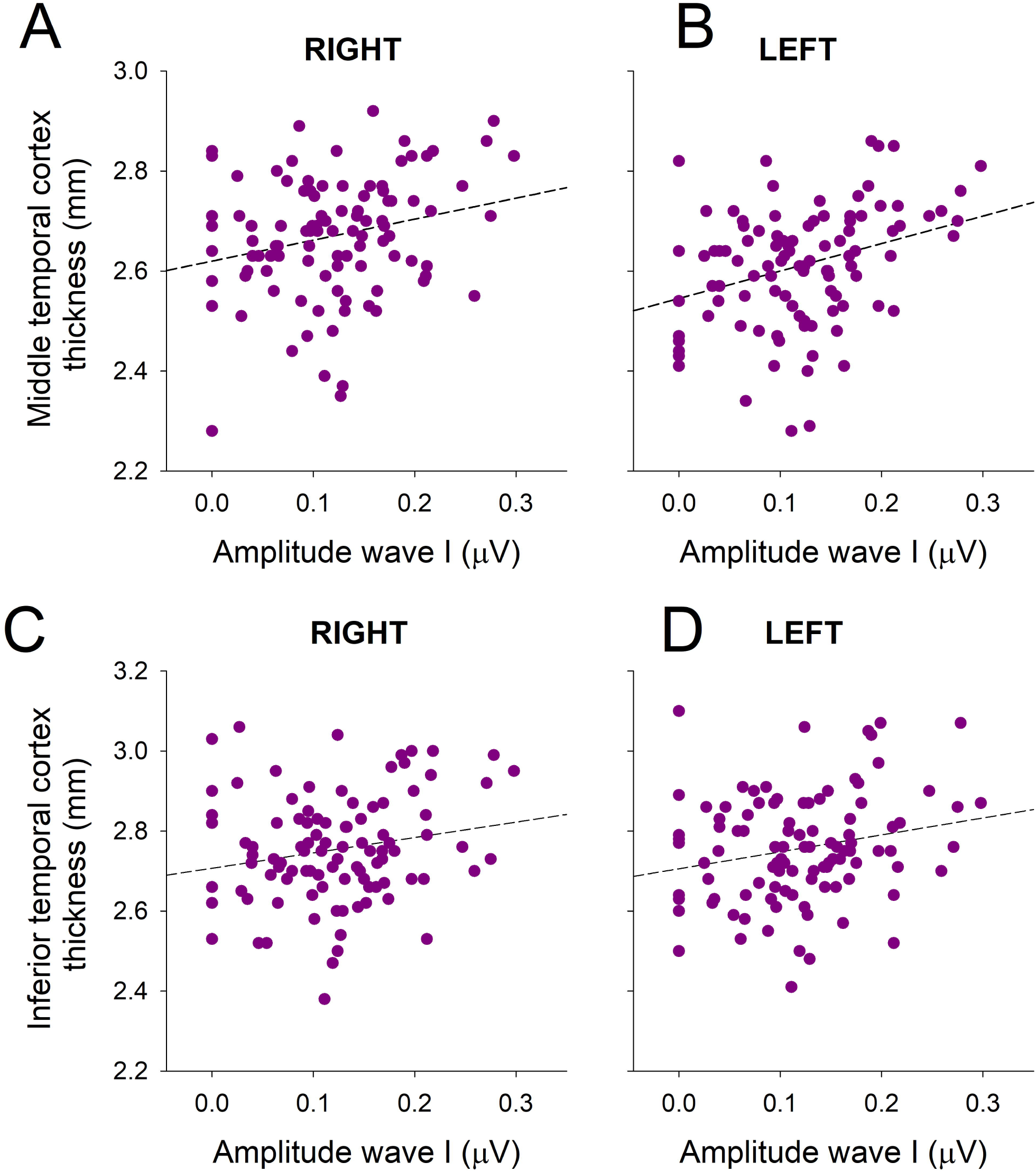
The thickness of bilateral middle and inferior temporal cortex is correlated with the amplitude of ABR wave I responses. (A) Right (rho=0.221; p=0.029) and (B) left (rho=0.263; p=0.009) middle temporal thickness correlated with wave I amplitude. (C) Right (rho=0.240; p=0.018) and (D) left (rho=0.216; p=0.034) inferior temporal cortex thickness correlated with wave I amplitude.

## Discussion

Here we give evidence that a reduced amplitude of suprathreshold auditory nerve responses (wave I) is associated with slower processing speed (TMT-A, digit symbol) and thinner bilateral temporal cortex in non-demented elderly humans. In addition, we show that the wave V/I ratio as a function of wave I amplitude yielded an asymmetric distribution, suggesting a midbrain compensatory gain increase for reduced suprathreshold auditory nerve responses.

### Aging, audiogram thresholds and suprathreshold ABRs

Although, in our data we did not find any significant correlation between the suprathreshold amplitudes of waves I and V with age (Figure 1), these results should be taken carefully, as the range of age of our subjects was between 65 and 85 years, and probably if we extend the range of age to younger subjects, it is very likely that we would find significant age effects. Indeed, previous studies performed in animals [33,34] as well as in humans [35-37] found significant reductions in wave I amplitudes with age.

In our study we also found that the amplitudes of suprathreshold ABR responses were not associated with audiogram thresholds (PTA calculated between 0.5 and 4 kHz), suggesting that auditory thresholds and suprathreshold functions are independent measures of auditory processing. In this line, we previously showed that a deteriorated hearing threshold function as evidenced by a reduced number of DPOAE is associated with atrophy of the anterior cingulate cortex and executive dysfunction in presbycusis [9]. In contrast, here we show that a reduced amplitude of suprathreshold auditory nerve responses is not associated with deteriorated executive function, but with slower processing speed (longer TMT-A latencies and worse digit symbol scores) and thinner temporal cortex. These findings suggest that the impairment of different auditory functions (threshold and suprathreshold) could affect different brain structures and cognitive domains.

### Midbrain gain increase

We found an increased wave V/I ratio in the group with reduced suprathreshold auditory nerve responses (<0.15 μV), which was independent of age and hearing thresholds. The gain increase of midbrain responses is also supported by the fact that the amplitudes of wave V responses were similar between the two groups with different wave I amplitudes (Table 2). Thus, the preserved amplitude of suprathreshold wave V responses in the group with reduced wave I could be reflecting a compensatory gain increase in the midbrain. A similar mechanism has been proposed for peripheral de-afferentation [16,38]. Moreover, animal models have shown that cochlear de-afferentation is sufficient for inducing an increase in the spontaneous activity of auditory cortex neurons [39], showing that the effects of peripheral de-afferentation can also affect cortical processing. Here we show in humans, that the group with reduced auditory nerve amplitudes has structural brain changes that were located bilaterally in the middle and inferior temporal cortex, and in the posterior cingulate cortex of the right hemisphere.

### Brain atrophy in presbycusis

Previous studies have related audiogram threshold loss with right temporal and cingulate cortex atrophy [9-12,40,41]. Here we extended these results, showing that in addition to audiogram threshold elevation, reduced suprathreshold amplitudes of auditory nerve responses are associated to a reduction of the thickness of bilateral middle and inferior temporal cortices, and right posterior cingulate cortex. Importantly, in the present study, we showed significant reductions in the cortical thickness, but not in the cortical volume of these regions. These results suggest that the cortical thickness is a more sensitive measure than cortical volume loss for evidencing brain atrophy related to suprathreshold auditory impairments. In addition, our data show that these structural brain changes can be present in earlier stages of presbycusis, or even in subjects with normal hearing (at least as evaluated by audiogram thresholds between 0.5 and 4 kHz).

In a previous work [9], we demonstrated that reduced PCC thickness was correlated with worse auditory thresholds in patients with presbycusis and cochlear dysfunction, suggesting that the atrophy of the right PCC is related to hearing loss. Here, we showed that a reduction in the cortical thickness of the right PCC is also associated with suprathreshold hearing impairments, suggesting that PCC atrophy is related to hearing threshold and suprathreshold impairments. The right posterior cingulate cortex is important for visuospatial abilities like orientation and spatial navigation. Interestingly the PCC is among the earliest regions that get atrophied in prodromal and preclinical Alzheimer’s disease [42]. In this line, the right PCC might be an important brain region linking hearing impairments with cognitive decline in presbycusis.

### Processing speed and suprathreshold auditory-nerve function

Previous evidence has shown that worse audiogram thresholds [10,43] or an alteration of the cochlear function as evidenced by loss of DPOAE [9] are associated with executive dysfunction, memory loss and global cognitive decline. In addition to these associations, here we show that reduced suprathreshold auditory-nerve responses are associated to slower processing speed, as evidenced by TMT-A responses (Figure 3, Table 3) and digit symbol performance (Table 3), cognitive tests which do not rely on auditory inputs. Processing speed tests are usually categorized as “fluid cognition” and are influenced by the aging process, but also by sensory impairments [44]. One speculative explanation for the association between reduced amplitude of auditory-nerve responses and slower processing speed could be related to the physiological aging process, resulting in loss of synapses at different levels of the nervous system [45]. In this sense, we can propose that due to the aging process, the loss of synapses between the inner hair cells and auditory nerve neurons would result in reduced amplitude of suprathreshold wave I responses [19], while reduced synapses at the central nervous system would lead to slower processing speed [45]. Although cochlear synaptopathy has been associated to loss of synapses due to acoustic trauma, it could also be an indirect measure of a general loss of synapses in the central nervous system, and therefore the greater the loss of synapses in different circuits of the nervous system, the slower is the processing speed. Another speculative explanation is that processing speed could be related to white matter microstructural changes in the peripheral and central auditory pathways, including the auditory nerve, as a reduced fractional anisotropy in diffusion tensor imaging has been demonstrated in diverse white matter tracts of patients with hearing loss [46].

## Conclusion

We conclude that a reduction of the suprathreshold amplitude of auditory nerve responses is related to slower processing speed and reduced cortical thickness in bilateral middle and inferior temporal cortices and in the right posterior cingulate cortex. Taken together, the present and our previous findings [9] suggest that thresholds and suprathreshold hearing impairments are associated with different types of cognitive functions and brain structural changes.

## Acknowledgments

We thank Professor Luis Robles for his valuable comments. Funded by Proyecto Fondecyt 1161155 to P.H.D, Proyecto Anillo ACT1403, CONICYT BASAL FB008, Proyecto ICM P09-015F y Fundación Guillermo Puelma.

